# Ecological histories determine the success of social exploitation

**DOI:** 10.1101/2023.12.14.571652

**Authors:** Kaitlin A. Schaal, Pauline Manhes, Gregory J. Velicer

## Abstract

Ecological context often modifies biotic interactions, yet effects of ecological history are poorly understood. In experiments with the bacterium *Myxococcus xanthus*, resource-level histories of genotypes interacting during cooperative multicellular development were found to strongly regulate social fitness. Yet how developmental spore production responded to variation in resource-level histories between interactants differed greatly between cooperators and cheaters; relative-fitness advantages gained by cheating after high-resource growth were generally reduced or absent if one or both parties experienced low-resource growth. Low-resource growth also eliminated facultative exploitation in some pairwise mixes of cooperation-proficient natural isolates that occurs when both strains have grown under resource abundance. Our results contrast with previous studies in which cooperator fitness correlated positively with resource level and suggest that resource-level variation may be important in regulating whether exploitation of cooperators occurs in a natural context.

## Introduction

Individuals can gain in absolute fitness from interacting with conspecifics, a fitness response often referred to as social exploitation (Fiegna & Velicer 2005). Here we consider effects of ecological history on two types of social exploitation that differ in their evolutionary implications, cheating by behaviorally obligate defectors (Diggle *et al*. 2007; Harrison 2009; Velicer *et al*. 2000) and social exploitation by cooperation-proficient genotypes (Fiegna & Velicer 2005) (see *Terms in* Methods).

In cheating by behaviorally obligate defectors, individuals unable to express a cooperative trait (or that can express it only at a low level) exploit those expressing it at a high level to gain a relative fitness advantage over them (Diggle *et al*. 2007; Harrison 2009; Velicer *et al*. 2000). Cheating by behaviorally obligate defectors can reduce equilibrium levels of the cooperative trait in a population and might drive the trait to extinction (Doebeli & Hauert 2005; Fiegna & Velicer 2003; Frank 2003; Hamilton 1964; Lehmann & Keller 2006; Van Dyken *et al*. 2011). Evolutionarily obligate cheaters rely on exploitation of cooperators to persist (Fiegna & Velicer 2003; Velicer & Vos 2009).

In contrast, evolutionarily facultative exploitation does not threaten the persistence of cooperation *per se* because in this case all interactants are genetically capable of expressing the cooperative trait (Buttery *et al*. 2009; Fiegna & Velicer 2005; Nair *et al*. 2018; Strassmann *et al*. 2000). One behavioral mechanism that might result in such exploitation is modulating the degree to which a cooperative trait is expressed as a function of social context. For example, a behaviorally facultative defector might express the cooperative trait at a high level when interacting with clonemates or other close kin but reduce its expression when interacting with less related individuals (Buttery *et al*. 2009). Evolutionarily facultative exploitation can alter the relative frequencies of different cooperative genotypes in a population without lowering the overall frequency of genotypes capable of cooperation.

Microbes exhibit diverse forms of cooperation (Branda *et al*. 2001; Crespi 2001; Diggle *et al*. 2007; Fiegna *et al*. 2023; Griffin *et al*. 2004; Harrison & Buckling 2009; Manhes & Velicer 2011; McGrath *et al*. 2004; Pande *et al*. 2020; Pesci *et al*. 1997; Rainey & Rainey 2003; Ross-Gillespie *et al*. 2009; Sandoz *et al*. 2007; Shapiro 1998; Strassmann *et al*. 2000; Velicer *et al*. 2000; Wilder *et al*. 2011; Zusman *et al*. 2007), but ecological factors can influence the relative fitness of cooperators and others that benefit from interacting with them, such as cheaters (Brockhurst *et al*. 2008, 2011; Harrison *et al*. 2008; Kümmerli *et al*. 2009b). For example, environmental toxins such as antibiotics (Vasse *et al*. 2017) or heavy metals (O’Brien *et al*. 2014) can benefit cheaters, while oxidative stress (García-Contreras *et al*. 2015) and viscosity (Kümmerli *et al*. 2009a) can benefit cooperators. High diversity (O’Brien *et al*. 2022), competitors (Harrison *et al*. 2008), or predators (Friman *et al*. 2013) within communities can also benefit cooperators. In some cases, phage presence benefits cooperators (Morgan *et al*. 2012; Saucedo-Mora *et al*. 2017), but in others it benefits cheaters (O’Brien *et al*. 2019; Vasse *et al*. 2015).

Additionally, both nutrient identity (Sexton & Schuster 2017) and level (Brockhurst *et al*. 2008) during cooperation can impact cooperation’s benefits and costs (Fronhofer *et al*. 2018). For example, experiments with bacteria have tested for nutrient-level effects on cooperator and cheater fitness during cooperation mediated by diffusible public-good molecules. High resource availability during cooperation was found to increase cooperator fitness - apparently by decreasing the relative cost of cooperation compared to low-resource conditions - although cheating was not eliminated (Brockhurst *et al*. 2008).

However, the fitness of cooperators relative to defectors may also be impacted by prior ecological conditions. Ecological histories are known to influence a variety of inter-specific interactions (Grigaltchik *et al*. 2012; Jones & Hurst 2023; Mcghee *et al*. 2012) and may also influence the social fitness of microbes. The plausibility of this hypothesis is illustrated by experiments with the social amoeba *Dictyostelium discoideum*, which develops into spore-bearing fruiting bodies in response to starvation. *D. discoideum* cells can differ in their propensity to form spores during development due to variations in their nutritional histories (“Chapter 9: Differentiation and Adhesion in the Aggregate” 2001; “Chapter 10: The Behavior of Cells in the Slug” 2001; Dubravcic *et al*. 2014; Forget *et al*. 2021; Kuzdzal-Fick *et al*. 2010). For example, during starvation, *D. discoideum* previously grown in glucose media form proportionally more spores than cells grown without glucose (Castillo *et al*. 2011; “Chapter 9: Differentiation and Adhesion in the Aggregate” 2001; “Chapter 10: The Behavior of Cells in the Slug” 2001; Leach *et al*. 1973). If co-developing defectors and cooperators were to differ in their nutritional histories, this would therefore impact their relative fitness. Nutritional histories of co-developing cooperator and defector cells might also matter for their relative fitness during development if they differ genetically in how nutrient-history variation impacts their developmental performance.

We investigate the effect of nutritional history on social exploitation in the soil bacterium *Myxococcus xanthus*, which forms spore-filled fruiting bodies upon starvation through a developmental process involving the exchange of several intercellular signals (Kroos 2017; Yang & Higgs 2014). Groups of tens-of-thousands of *M. xanthus* cells aggregate, but only a small minority become spherical spores; the remaining cells either die or remain rod-shaped cells (Amherd *et al*. 2018; Lee *et al*. 2012; Zusman *et al*. 2007). In this system, developmental cheaters are defined as strains that are defective at spore production in monoculture (compared to high-sporulating cooperators) but which produce proportionally more spores than cooperators in mixed groups (Schaal *et al*. 2022; Velicer *et al*. 2000). Laboratory experiments have repeatedly shown that developmental cheaters can rapidly increase in frequency due to their exploitation of cooperators, in some cases leading to large population crashes and even extinction events (Fiegna & Velicer 2003; Manhes & Velicer 2011; Velicer *et al*. 2000). Facultative exploitation during development has previously been observed between *M. xanthus* natural isolates that all sporulate at similarly high levels in monoculture (Fiegna & Velicer 2005; Vos & Velicer 2009).

We tested for the effects of nutritional history on both cheating and facultative exploitation. We cultured a cooperator strain and multiple cheater strains of *M. xanthus* in both high- and low-resource growth media before mixing the cooperator with each cheater genotype pairwise in all possible nutrient-history combinations and then quantifying spore production by each genotype after starvation-induced development. By examining multiple cheaters, we tested both i) whether nutrient-history variation impacts cooperator-cheater interactions, including potentially determining whether cheating occurs or not, and ii) whether any such nutrient-history effects are similar or different across distinct cheater-cooperator genotype pairs. Similarly, we examined the effects of nutritional history on relative developmental fitness in pairwise mixes of natural isolates with only short histories of lab cultivation that all sporulate at a high level in monoculture. Each of these natural isolates had previously been shown to either exploit or be subject to exploitation by another natural isolate during development (Fiegna & Velicer 2005).

## Materials and Methods

### Terms

The definitions of ‘cheating,’ ‘defection,’ and ‘exploitation’ (and their variants) adopted here are based on overall patterns of quantitative phenotypes. In this study, the core phenotype is how many spores strains produce across multiple social contexts: i) comparison of distinct genotypes across clonal populations (in the case of defection), ii) comparison of a single genotype between clonal and mixed populations (exploitation), or iii) both comparison of individual genotypes across clonal vs mixed populations and comparison of distinct genotypes within mixed populations (cheating). Fulfillment of the criteria for these definitions does not imply any degree of adaptation for increased proficiency at the observed behavior, *e*.*g*. adaptedness for exploitation or cheating. The questions of whether, in what manner, and to what degree an observed exploitative behavior evolved adaptively are distinct from the question of whether the behavior is observed in any particular context in the first place; such evolutionary questions require separate criteria and investigation. Additionally, the occurrence of exploitation, defection, or cheating during social interactions does not imply that the absolute fitness of cooperators is reduced by such interactions, although such reductions will often be expected, for example when behaviorally obligate defectors present at a relatively high frequency in a population cheat on cooperators and thereby reduce total group productivity (Fiegna & Velicer 2003; Velicer *et al*. 2000; Velicer & Vos 2009).

### Bacterial strains and culturing

In the following assays, we examined pairwise social interactions among several lab strains and, separately, among several natural isolates of *M. xanthus*.

#### Lab strains

As the sporulation-proficient cooperator, we used *M. xanthus* strain GJV1 ((Velicer *et al*. 2006); strain S in (Velicer *et al*. 1998), a derivative of DK1622 that is hereafter referred to as WT for ‘wild-type’). As obligately defecting cheaters, we used two strains derived from the MyxoEE-1 evolution experiment (Rendueles & Velicer 2020): GJV9 (a rifampicin-resistant variant of GVB207.3 (Manhes & Velicer 2011)) and GVB206.3 (also rifampicin resistant (Velicer *et al*. 2000)). Here, we refer to these cheaters as Ch1 and Ch2, respectively. Each was isolated from independently evolved populations descended from GJV1 after 1000 generations of growth in nutrient-rich liquid medium. Their cheating behaviors have been repeatedly demonstrated (Fiegna *et al*. 2006; Fiegna & Velicer 2003; Velicer *et al*. 2000; Velicer & Stredwick 2002).

#### Natural isolates

For interactions between less-domesticated *M. xanthus* natural isolates, we used previously described antibiotic-resistant mutants of isolates DK801 (kanamycin-resistant) and Mxx41 (novobiocin-resistant) and the isolate Mxx144 (sensitive to both antibiotics), all of which sporulate at similar levels in monoculture (Fiegna & Velicer 2005). In the study that first documented social exploitation during sporulation among these strains (Fiegna & Velicer 2005), DK801 and its mutant were collectively referred to as strain D, Mxx41 and its mutant were collectively referred to as strain G, and Mxx144 was referred to as strain I. For consistency with (Fiegna & Velicer 2005), hereafter in this study we still refer to the resistant mutants of DK801 and Mxx41 as strains D and G, respectively, and Mxx144 as strain I.

#### Growth conditions

In the following assays, we grew strains in either high-nutrient or low-nutrient conditions before inducing development. The high-nutrient condition (H) was CTT liquid (Hodgkin & Kaiser 1977), which contains 1% Casitone. The low-nutrient media (L) had the same composition as CTT except with reduced Casitone (0.1% for lab strains and 0.2% for natural isolates). We grew the bacteria in liquid at 32 °C and either 300 rpm (lab strains) or 200 rpm (natural isolates) and maintained them in exponential phase until starting the experiment.

### Developmental assays

We grew all strains in high- and low-nutrient liquid media (Fig. 1), then centrifuged them at 5000x*g* for 15 minutes at 20 °C. We resuspended the pellets in TPM buffer (Kuspa *et al*. 1986) to a density of ∼5 x 10^9^ cells/ml. For pure-culture assays, we spotted 100 µl of each culture onto TPM 1.5% agar plates. For lab-strain mixed-culture assays, we combined the cheater strain with the WT at a 1:9 ratio; for natural isolate mixes, we used a 1:1 ratio as in (Fiegna & Velicer 2005). We spotted the entire 100 µl of each mixture onto TPM 1.5% agar plates. We incubated all plates at 32 °C and 90% rH for 5 days before harvesting each population into ddH_2_O, heating for 2 hours at 50 °C to kill non-spore cells, and sonicating to disperse spores. We plated various dilutions of the spores into CTT 0.5% agar and incubated at 32 °C and 90% rH until colonies were visible. For lab strains, we plated samples in agar with and without 5 µg/ml rifampicin to obtain counts of cheater spores and total spores, respectively. For natural isolate mixes, we counted colonies of strain D in agar with 40 µg/ml kanamycin and colonies of strain G in agar with 10 µg/ml novobiocin; we estimated spore production by strain I in mixes by subtraction of selective-agar colony counts from non-selective-agar colony counts. When we counted no spores at the lower limit of detection, we estimated 0.9 spores. We performed four temporally separate replicates for all assays.

**Figure 1.**
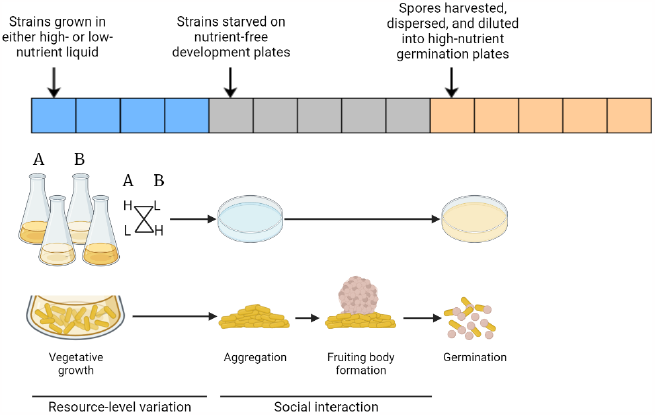
Resource history and experimental design. Here we show a timeline of the resource abundance experienced by the bacterial populations throughout the experiment, the methodological steps, and the respective life cycle stages of the *Myxococcus xanthus* populations. Each segment of the timeline represents one day, and the color of the square represents the resource abundance (blue = variable depending on treatment, grey = absent, orange = high). The flasks show strains A and B being grown under either high- or low-resource conditions and then being mixed pairwise under all resource-history combinations before plating onto nutrient-free plates.

Using pure-culture sporulation assays, we tested whether the use of antibiotic as a selection agent alters the number of spores produced by D or G that germinate and grow into visible colonies (difference of means +/- antibiotic [95% confidence intervals], n = 4 for all; D_high_ = 0.31 [-0.37, 0.99], D_low_ = 0.18 [-0.65, 1.02], G_high_ = 0.01 [-0.20, 0.23], G_low_ = 0.07 [-0.27, 0.41]; we considered this reasonable evidence for equivalence and proceeded.

### Measures of cheating and exploitation

#### Cheating by lab strains

We tested for the ability of each known cheater strain to cheat on WT during sporulation for all four combinations of pre-development nutritional histories. We assessed cheating by calculating the relative fitness parameter *W*_*ij*_ (Manhes *et al*. 2022; Schaal *et al*. 2022):

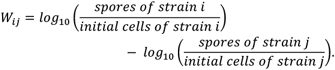

Positive and negative values of this parameter indicate that the cheater exhibits higher or lower relative spore production than WT in pairwise mixes.

#### Mixing effects among natural isolates

Following (Fiegna & Velicer 2005), we estimated for each strain the mixing effect parameter *C*_*i*_*(j)* which compares the sporulation efficiency of a focal strain *i* in a pairwise-mixed competition with strain *j* relative to the sporulation efficiency of strain *i* in pure culture. The sporulation efficiency *D*_*i*_ of strain *i* in pure culture is the ratio of cells that survive as viable spores:

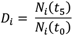

*N*_*i*_(*t*_*5*_) is the viable population size after 5 days of development and *N*_*i*_(*t*_*0*_) is the viable population size before development, here 5 x 10^8^ cells.

The sporulation efficiency of strain *i* when mixed with strain *j* is given by

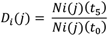

where *N*_*i*_(*t*_*0*_) is 2.5 x 10^8^ cells.

The effect of strain *j* on the sporulation of strain *i* when the two strains are mixed relative to sporulation by strain *i* in pure culture is given by

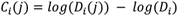

Positive and negative values of C_*i*_(*j*) indicate that strain *i* generates proportionally more or fewer spores when mixed with strain *j* than when alone, respectively. Mixing-effect values were calculated to test whether growth conditions affected mixing outcomes.

### Data analysis

We analyzed our data in R version 4.3.0 (R Core Team 2018) and RStudio version 2023.03.0 (RStudio Team 2020) using the packages “tidyr” (Wickham *et al*. 2023b), “dplyr” (Wickham *et al*. 2023a), and “DescTools” (Signorell 2023), and we visualized them using the “ggplot2” package (Wickham 2016). We tested the effect of nutritional history using generalized linear models (GLMs) followed by *t*-tests or Tukey HSD tests (see *Results*). Original data and analysis scripts can be found at https://github.com/kaschaal/resource-scarcity-cheating. For clarity, we generally present statistical results in tables rather than referenced in-line.

## Results

### Low-nutrient resource histories can prevent cheating

To examine effects of prior resource levels on the fitness of obligate defectors while interacting with a cooperator, we compared relative sporulation by the two cheaters when mixed pairwise with WT after growth in liquid culture with variable nutrient concentrations. We first tested for potential effects of resource level during growth on subsequent starvation-induced spore production in pure cultures of WT and the two cheaters. For all three strains, no effect of nutritional history was detected (Fig. 2A; Table S1 Test 1).

**Figure 2.**
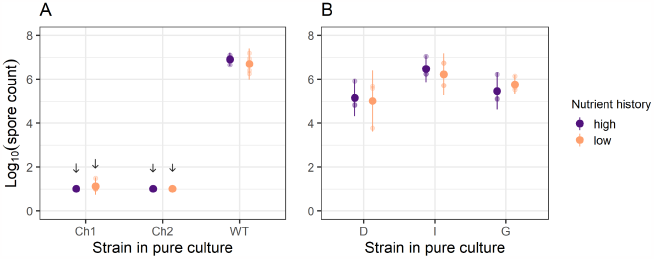
Resource-level history does not alter spore production of single-genotype populations. Total spore production in pure culture by two cheaters and a cooperator reference strain (A) and three natural isolates (B) used in this study after prior growth at high- or low-nutrient conditions. Small dots show individual-replicate estimates and large dots show cross-replicate means; 95% confidence intervals are represented. No strain showed a significant difference in sporulation ability between high vs low nutrient-level histories (Table S2, Test 1). Downward arrows indicate treatments in which zero spores were detected at the lower limit of detection (see *Methods*) in at least one replicate.

We then tested for effects of nutritional history on cheater vs. WT fitness outcomes in chimeric groups. Each cheater was mixed with WT in a 1:9 ratio in all four possible resource-history combinations (H/H, H/L, L/H, L/L). For each strain-pair and resource-history combination, we calculated the sporulation fitness of the respective cheater genotype relative to that of WT in the same mix, as reflected by the parameter *W*_*ij*_ (see *Methods*). In these calculations, genotype *i* represents the cheater and genotype *j* represents WT. ANOVAs revealed significant generic effects of WT nutrient history on *W*_Ch1:WT_ and of both WT and Ch2 nutrient histories on *W*_Ch2:WT_ (Table S1, Tests 2-3). Thus, resource history impacted social fitness during intergenotype interactions without having altered the performance of individual genotypes in isolation.

Following from the ANOVA results, we tested whether particular nutrient-history combinations determined whether cheating occurred or not. We say that cheating occurred in cases where *W*_*Ch:WT*_ is significantly greater than zero (see *Methods*), *i*.*e*., the cheater produces proportionally more spores than WT in co-culture. In accordance with previous studies, both cheaters exhibited strong cheating when both the cheaters and WT had grown under standard lab conditions (H/H), as reflected by significantly positive values of *W*_*Ch1-H:WT-H*_ and *W*_*Ch2-H:WT-H*_ (Fig. 3A; Table S2 Tests 4 & 5).

**Figure 3.**
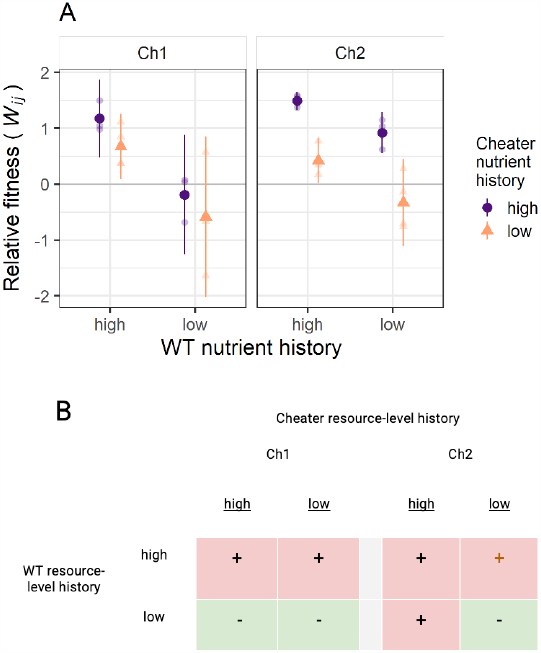
Historical resource scarcity eliminates cheating in some strain-history combinations. (A) Relative sporulation outcomes in pairwise mixes of each cheater strain with WT (*W*_*ij*_, *i =* cheater; *j =* WT) for all four possible resource-level-history combinations for each strain pair. Small dots show individual-replicate estimates and large dots show cross-replicate means; error bars show 95% confidence intervals. (B) Qualitative cheating-occurrence outcomes; whether cheating occurred (+) or not (-) is shown for all strain-pair/growth-history combinations. The brown + symbol indicates a case in which cheating occurred but cheater fitness was significantly lower than when both strains had grown at high resource levels.

We found that Ch1 was able to cheat whenever WT had grown under high-resource conditions, irrespective of Ch1’s own growth history (Fig. 3A; Table S2 Tests 4 & 6 [*p* = 0.04]). However, Ch1 failed to cheat whenever WT had experienced low-nutrient conditions, again irrespective of Ch1 history (Fig. 3A; Table S2 Test 6). For Ch2, in addition to cheating under H/H conditions, cheating occurred if only Ch2 or WT had experienced low-nutrient conditions, but was not observed when both had experienced low-nutrient conditions (Fig. 3A; Table S2 Test 6).

In other words, a history of low-nutrient growth for WT eliminated cheating by Ch1 regardless of Ch1’s nutritional history. Low-nutrient growth by WT also eliminated cheating by Ch2, but only when Ch2 had also experienced low-nutrient growth (Fig. 3B). Thus, the ecological history of a cooperator determined whether it was susceptible to cheating by two distinct cheaters, although exactly when WT history determined the occurrence of cheating (with respect to cheater history) differed across the two cheater genotypes.

### Low-nutrient history can reduce cheater performance even when cheating still occurs

We further analyzed the effects of resource history on cheater performance by testing for differences between *W*_*ij*_ under H/H conditions versus under the non-standard conditions. Growing only the cheater with low nutrients had no effect on Ch1 fitness compared to H/H histories (Fig. 3A; Table S3 Test 7), but it reduced relative fitness for Ch2 (Table S3 Test 8) without eliminating cheating (Table S2 Test 6, *p* = 0.044).

In general, cheater relative fitness was reduced for both cheaters when WT was grown under low-nutrient conditions compared to WT growth at high resource levels (Figs. 3A, S1; Table S1 Tests 2 & 3). However, when only WT was grown with low nutrients, Ch2 performance may have been reduced relative to the Ch2-H/WT-H treatment, but the result was not significant (Table S3 Test 8), and Ch2 still cheated (Table S2 Test 6, *p* = 0.012). We found similar results for Ch1 (Table S3 Test 7), although in this case we found no evidence of cheating (Table S2 Test 6, *p* = 0.86).

When both strains were grown with low nutrients, cheater performance was significantly reduced (Table S3 Tests 7 & 8). For Ch1, we don’t see a significant difference between Ch1-L/WT-L and either Ch1-H/WT-L (Table S3 Test 7) or Ch1-L/WT-H (Table S3 Test 7). However, for Ch2, there is a significant difference between Ch2-L/WT-L and both Ch2-H/WT-L (Table S3 Test 8) and Ch2-L/WT-H (Table S3 Test 8).

To summarize, low-nutrient growth by one or both partners reduced cheater performance in some cases. For Ch1, this result is clear for the L/L condition, and there is the suggestion of an effect for Ch1-H/WT-L. There is evidence that Ch2 performance is reduced when either or both of the interacting strains has a prior history of lower nutrients, although the result for Ch2-H/WT-L is not quite significant; for Ch2-L/WT-H cheater performance is significantly reduced but cheating still occurs.

### Low-nutrient growth can prevent subsequent exploitation between natural isolates

To test for effects of nutrient availability during growth on exploitative relationships among three natural isolates during development, we compared the outcomes of pairwise competitions between the sporulation-proficient isolates for all four possible growth-history combinations (H/H, H/L, L/H, L/L). We calculated the mixing-effect parameter *C*_*i*_(*j*) (see *Methods*), where a positive value indicates exploitation of strain *j* by *i*. Irrespective of their nutrient-level history, all three strains sporulate at high levels in pure culture (Fig. 2), but strain I produces more spores than strains D and G (Fig. 2B; Table S4 Tests 9 & 10). As above, we tested for potential effects of nutritional history on subsequent developmental spore production in pure cultures. For all strains, no effect of nutrient level during growth was detected (Fig. 2B; Table S4 Test 11).

After growth under standard lab conditions (H/H), patterns of both relative spore production and social exploitation were generally similar to the outcomes of developmental competitions with these same strains conducted after H/H growth in a previous study (Fiegna & Velicer 2005), which demonstrated a clear fitness hierarchy (D > I > G). We again find clear evidence of strain G’s low relative fitness (Figs. 4, S2); while strain D appears to have produced more spores than strain I in the H/H D:I mix, high variability in strain I spore estimates across replicates makes this unclear. Contrary to previous observation (Fiegna & Velicer 2005), apparent exploitation of strain I by strain D after H/H growth was not significant in our experiments (Fig. 4; Table S4 Test 12). However, consistent with previous observations (Fiegna & Velicer 2005), strains D and I both exploited strain G (Fig. 4; Table S4 Tests 13 & 14).

**Figure 4.**
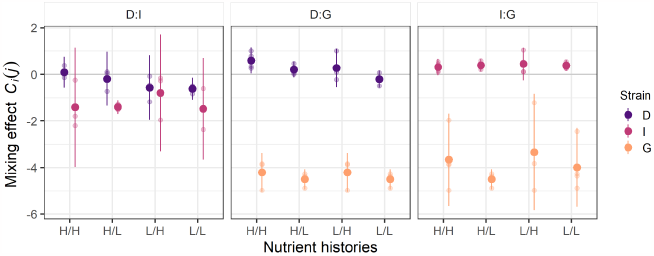
Historical resource scarcity can reduce or eliminate exploitation between distinct cooperators. The effects of pairwise mixing of two cooperation-proficient natural isolates on the sporulation efficiency of each paired strain - *C*_*i*_*(j)* (see Methods) - are shown for all four possible growth-history combinations for each of three strain pairs: (A) strains D vs G; (B) strains D vs I; (C) strains I vs G. *x*-axis labels identify the growth history of each partner (H = high-nutrient history, L = low-nutrient history). The first- and second-listed nutrient-history indicators refer to the first- and second-listed strains of each pair. The first-listed strain of each pair is the strain previously shown to exploit the other after growth under standard lab conditions (H/H). The absence of any mixing effect on a given strain would correspond to a *C*_*i*_*(j)* value of 0; exploitative and antagonized responses to mixing are indicated by significantly positive and negative *C*_*i*_*(j)* values, respectively. Small dots show individual-replicate estimates and large dots show cross-replicate means; error bars represent 95% confidence intervals.

Focusing first on the competitions between D and I, we found no evidence of D exploiting I in any of the three treatments in which at least one strain had been grown at the low resource level; all three *C*_*D*_(*I*) estimates were negative (Fig. 4). There was even a suggestion that D might be harmed in mixture with I under L/L conditions, although the result is not significant (Table S4 Test 15). However, strain I was significantly harmed by interacting with D after D-H/I-L growth histories (Table S4 Test 15). We found no significant evidence for changes in *C*_*i*_(*j*) due to any specific nutritional-history combination (Table S4 Tests 16 & 17).

In competitions between D and G, spore production by D (the strong winner, Fig. 4) was sensitive to the nutritional status of both strains. Although D exploited G under H/H conditions (as reported above, Fig. 4), strain D failed to show clear evidence of exploitation if one strain, but not the other, was grown under low resources (Table S4 Test 18). Strikingly, exploitation by D clearly did not occur after both had been grown at low nutrient levels (Table S4 Test 19). Spore production by G in mixture was extremely low, being reduced by a factor >10^4^ (and to less than 0.01% of total spores; Fig. S2) under all conditions.

In mixes of strains I and G, mean *C*_*I*_(*G*) estimates suggest that strain I might exploit strain G in all growth-history treatments. *C*_*I*_(*G*) was significantly positive in three of the four treatments (Fig. 4; Table S4 Tests 13, 14 & 21), and in both treatments in which strain G had grown in low-nutrient medium (H/L and L/L; Table S4 Test 21), but not when strain I had grown under low nutrients and strain G had grown under high nutrients (Table S4 Test 21, *p*_I-L/G-H_ = 0.14). As in mixtures with strain D, G sporulated poorly in all treatments, consistently accounting for less than 0.1% of the total spore count (Fig. S2).

Although details differed, in several of the natural-isolate mixes, low-nutrient growth by either or both competitors reduced (or appeared to reduce) spore production by the dominant competitor relative to the respective H/H pairings. Thus, for some genotype combinations, low-nutrient growth before development in chimeric groups is found to prevent not only cheating by sporulation-defective strains (Fig. 3), but also social exploitation by sporulation-proficient natural isolates (Fig. 4).

## Discussion

### Resource history determines whether cheating or exploitation happens

Both in pairs of a cooperation-proficient lab reference strain of *M. xanthus* and multiple cheater strains, and in pairs of natural isolates with only short histories of lab cultivation, the pre-development resource-level history of co-developing strains is found to strongly affect social fitness during sporulation. An abiotic ecological gradient changes the character of exploitative social interactions in microbes, just as abiotic gradients can alter interspecific interactions (Friede *et al*. 2016; Johnson *et al*. 1997; Jones & Hurst 2023). For three of five strain pairs examined, low-resource growth by one or both strains in the pair eliminated a clear exploitative interaction observed when both strains were grown with high resources. In one pairing with a cheater, low-nutrient growth by either party sufficed to prevent cheating, whereas in the other pair, cheating was only eliminated when both strains grew with low nutrients (Fig. 3). In one pair of natural isolates (D and G), low-nutrient growth by either competitor was sufficient to remove clear evidence of exploitation (Fig. 4).

### Mechanism hypotheses

Resource level during growth altered developmental competition outcomes by differentially impacting strain interactions rather than the intrinsic sporulation abilities of paired strains (Fig. 2). Several non-mutually-exclusive hypotheses might explain why low-resource growth decreased the sporulation efficiency of cheaters relative to cooperators. These include differential effects of growth history on (a) developmental timing, (b) signal response, and/or (c) cell lysis. Regarding timing, low-resource history might increase the time it takes cheater cells to differentiate into spores compared to cooperator cells (without impacting final total productivity), thereby allowing the cooperator to convert more of its cells into spores in mixed groups. The effect of low-nutrient conditions on sporulation would then be more pronounced in the early stages of development. This hypothesis could be tested by tracking temporal dynamics of spore production after growth in different media (Kuzdzal-Fick *et al*. 2010).

Regarding signaling, cheater cells in mixed groups with low–nutrient growth histories might be comparatively less responsive to developmental signals produced by the cooperator than they are in high-resource-history groups. This could manifest either in differences in sporulation timing (as outlined above) or in the conversion frequency of cheater cells into spores, irrespective of timing. Alternately, cooperators grown under low-resource conditions may not produce as many signal molecules as they do after high-resource growth, rendering them less cheatable. Regarding lysis, the comparative degree of cell lysis between strains might differ in high-vs low-resource-history groups.

### Implications for natural populations

Our results complicate the effort to understand cheating, and social exploitation more broadly, in natural populations of microbes. They suggest that estimating the frequency of genotypes in a natural population that exhibit cheating under one set of ecological conditions is unlikely to accurately reflect the overall prevalence of cheating in that population across space and over time.

*M. xanthus* is a predator of fungi and other bacteria (reviewed in (Thiery & Kaimer 2020)). Wild strains of *M. xanthus* certainly encounter diverse potential prey in their natural environments and likely often vary in their ability to kill and consume any given prey genotype (Morgan *et al*. 2010). Our results suggest that if some co-developing cells experienced a nutrient-poor prey environment just before aggregation, this may, in some cases, reduce or prevent cheating. Cheating may therefore be less common under some natural conditions than would be inferred from studies performed under standard laboratory conditions with nutrient-rich growth. We suggest testing outcomes of social conflicts during development after co-developing parties had grown on prey which differ in nutritional accessibility toward *M. xanthus*. Together, our results suggest that the occurrence and strength (or degree) of cheating in nature may be highly variable across different ecological contexts, including different prey environments.

### Resource-level effects on cooperation vary across social systems

Whatever the mechanisms at play, it is clear that previous findings that high resource levels reduce cheating due to reduced costs of cooperation (Brockhurst *et al*. 2008) do not reflect a universal rule of cooperative systems. Our results add to the body of work demonstrating that how distinct cooperative systems respond to environmental variables can differ (Ross-Gillespie *et al*. 2009; Van Dyken & Wade 2012), even directionally. Effects of resource level on cooperation need further investigation across diverse systems until a broad picture of the degree of variation in such effects emerges.

There are multiple differences between this study and an earlier study by Brockhurst *et al*. (Brockhurst *et al*. 2008) that might contribute to their contrasting outcomes. First, in the Brockhurst *et al*. experiments, growth fueled by manipulated resource levels occurred simultaneously with the expression of the focal cooperative behavior. In our experiments, the manipulated resource environments were experienced historically - before starvation, during which cooperation occurred. How resource-level variation relates to the costs and benefits of cooperation may differ as a function of the timing of such variation relative to when cooperation occurs.

Second, and relevant to temporal aspects, the cooperative trait of interest in our study – sporulation in the context of multicellular fruiting body development – is likely more mechanistically complex than the forms of public goods cooperation, such as siderophore production, that have been the focus of prior relevant studies. More complex forms of cooperation may be subject to a greater range of resource-level effects on interaction outcomes than simpler forms.

### Ecological history shapes social interaction

Species’ ecological histories can impact the outcomes of their interaction (Grigaltchik *et al*. 2012), in magnitude or even sign. For example, thermal histories can modify host-parasite-interaction phenotypes, such as when the degree to which *Spiroplasma* bacterial symbionts protect *Drosophila melanogaster* larvae from wasp attack is determined by the temperature at which the mother flies had previously developed, and not by the temperature at the time of wasp-larvae interaction (Jones & Hurst 2023). In predator-prey interactions between distinct species of bacteria, the very direction of predation can reverse as a function of the temperature at which one of the parties grew before the interaction (Vasse *et al*. 2023). Here, we see that prior resource histories of conspecifics interacting during a cooperative process can determine both i) whether cheating by genotypes obligately defective at cooperation occurs, and ii) whether social exploitation between cooperation-proficient competitors occurs. These findings encourage further investigation of how ecological histories shape social interactions between genetically distinct conspecifics across diverse species in which individuals may experience ecologically distinct environments before cooperating.

## Acknowledgements

We thank Nicola Mayrhofer for assistance with the literature search and Marco La Fortezza for helpful comments on the manuscript. This study was funded in part by by U.S. National Institutes of Health Grant GM079690, and SNF grants 31003A_160005 and 310030B_182830 to G.J.V.

## Supplemental Information for

**Figure S1.**
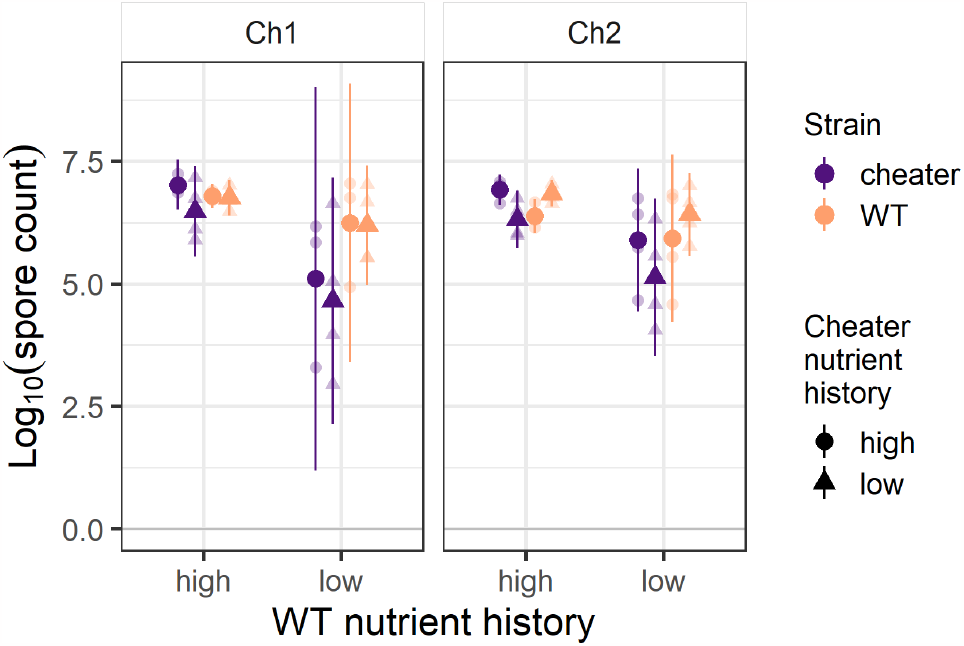
Absolute spore production by cheaters and WT in mixture. Spore production values by cheaters in mixture with WT used to calculate *W*_*ij*_ values in Figure 3. Small dots show individual-replicate estimates and large dots show cross-replicate means; error bars represent 95% confidence intervals.

**Figure S2.**
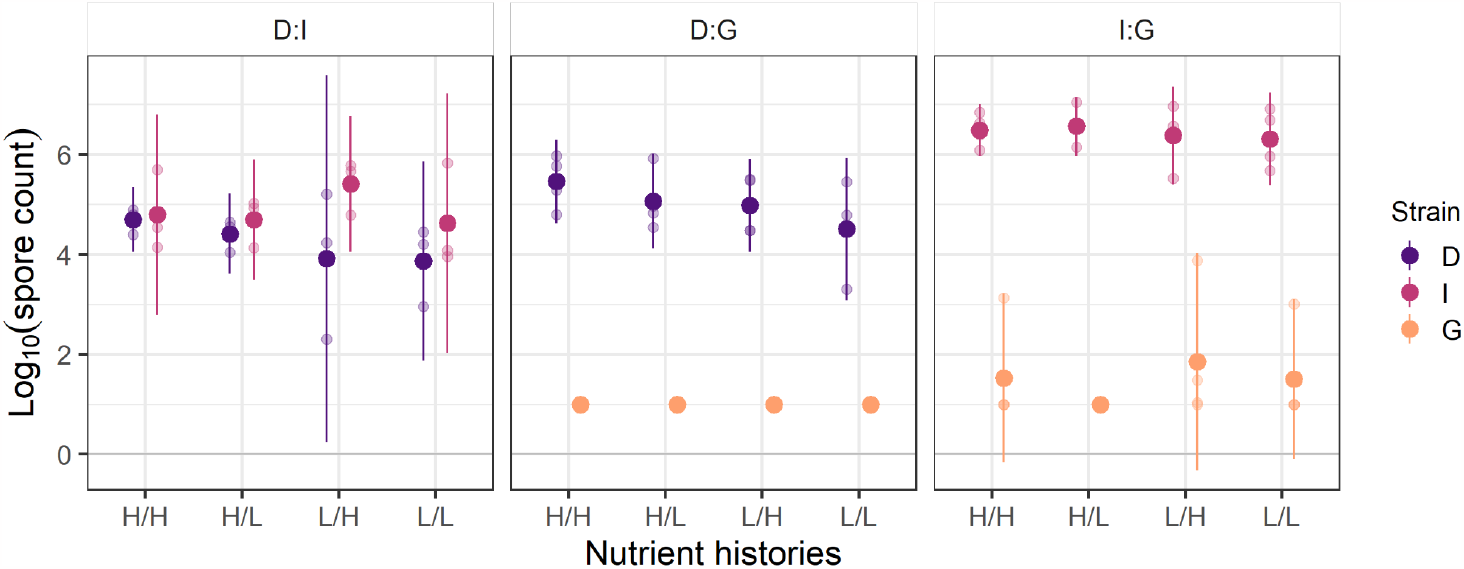
Spore production by natural isolates in mixture. Spore production values by natural isolates in pairwise mixes used to calculate *C*_*i*_*(j)* values for Figure 4. Spore counts for strain I were calculated by subtracting counts for either D or G from total counts obtained from nonselective dilution plates. Small dots show individual-replicate estimates and large dots show cross-replicate means; error bars represent 95% confidence intervals.

**Table S1.** Statistical tests for effects of nutrient histories (‘nutrients’) on pure-culture spore production (Test 1) and cheater *W*_*ij*_ values in competition with WT (Tests 2 and 3)

| # | Test | Variable(s) | F-statistic | p-value |
| --- | --- | --- | --- | --- |
| 1 | ANOVA for effect of strain x nutrients on $\log_{10}$(CFUs) | strain | $F_{2,17} = 1673.17$ | < 0.0001 |
| | | nutrients | $F_{1,17} = 0.18$ | 0.68 |
| | | strain x nutrients | $F_{2,17} = 0.95$ | 0.41 |
| 2 | ANOVA for effect of Ch1 nutrients x WT nutrients on $W_{ij}$ of Ch1 | nutrients[Ch1] | $F_{1,10} = 2.06$ | 0.18 |
| | | nutrients[WT] | $F_{1,10} = 17.56$ | 0.002 |
| | | nutrients[Ch1] x nutrients[WT] | $F_{1,10} = 0.03$ | 0.87 |
| 3 | ANOVA for effect of Ch2 nutrients x WT nutrients on $W_{ij}$ of Ch2 | nutrients[Ch2] | $F_{1,12} = 58.7$ | < 0.0001 |
| | | nutrients[WT] | $F_{1,12} = 19.2$ | 0.0009 |
| | | nutrients[Ch2] x nutrients[WT] | $F_{1,12} = 0.35$ | 0.56 |

**Table S2.** Statistical tests for whether cheating occurred in each strain-pair x nutrient-history combination.

| # | Test | Notes | t-statistic & df | p-value |
| --- | --- | --- | --- | --- |
| 4 | One Sample t-test of whether $W_{Ch1:WT} > 0$, H/H histories | Does cheating happen? A priori = yes | $t = 7.28$, df = 2 | 0.009 |
| 5 | One Sample t-test of whether $W_{Ch2:WT} > 0$, H/H histories | Does cheating happen? A priori = yes | $t = 29.75$, df = 3 | < 0.0001 |
| 6 | One Sample t-tests of whether $W_{Ch:WT} > 0$ for H/L, L/H and L/L histories; Bonferroni-Holm corrected | Ch1 H/L | $t = -0.75$, df = 2 | 0.86 |
| | | Ch1 L/H | $t = 3.64$, df = 3 | 0.04 |
| | | Ch1 L/L | $t = -1.3$, df = 3 | 0.86 |
| | | Ch2 H/L | $t = 8.05$, df = 3 | 0.01 |
| | | Ch2 L/H | $t = 3.36$, df = 3 | 0.04 |
| | | Ch2 L/L | $t = -1.34$, df = 3 | 0.86 |

**Table S3.** Statistical tests for differences in *W*_*ij*_ values between distinct nutrient-history combinations.

| # | Test | Comparison | difference in means [95% confidence interval] | p-value |
| --- | --- | --- | --- | --- |
| 7 | Post-hoc Tukey HSD tests from Test 2 ($W_{ij}$ of Ch1) | H-H - L/H | 0.5 [-0.86, 1.86] | 0.68 |
|  |  | H/H - H/L | 1.36 [-0.09, 2.82] | 0.07 |
|  |  | H/H - L/L | 1.76 [0.4, 3.12] | 0.01 |
|  |  | L/H - H/L | 0.86 [-0.5, 2.22] | 0.27 |
|  |  | L/H - L/L | 1.25 [-0.0006, 2.52] | 0.05 |
|  |  | H/L - L/L | 0.4 [-0.96, 1.76] | 0.81 |
| 8 | Post-hoc Tukey HSD tests from Test 3 ($W_{ij}$ of Ch2) | H-H - L/H | 1.07 [0.43, 1.71] | 0.002 |
|  |  | H/H - H/L | 0.57 [-0.06, 1.21] | 0.08 |
|  |  | H/H - L/L | 1.82 [1.19, 2.46] | < 0.0001 |
|  |  | L/H - H/L | 0.5 [-1.13, 0.13] | 0.15 |
|  |  | L/H - L/L | 0.75 [0.12, 1.39] | 0.02 |
|  |  | H/L - L/L | 1.25 [0.61, 1.89] | 0.0004 |

**Table S4.**
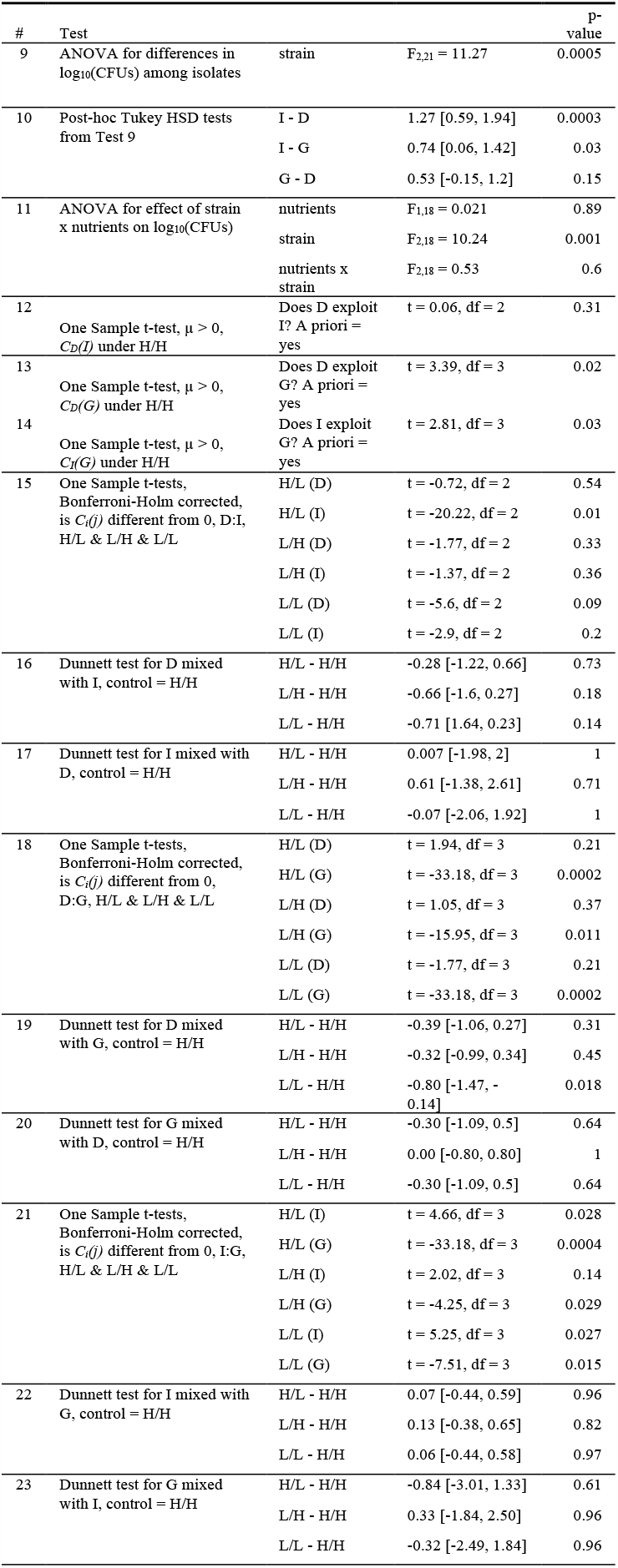
Statistical tests for effect of resource history on exploitation in natural isolates.

## Notes

https://github.com/kaschaal/resource-scarcity-cheating

### Competing Interest Statement

The authors have declared no competing interest.

